# A thermodynamic model of protein structure evolution explains empirical amino acid rate matrices

**DOI:** 10.1101/2020.12.02.408807

**Authors:** Christoffer Norn, Ingemar André, Douglas L. Theobald

## Abstract

Proteins evolve under a myriad of biophysical selection pressures that collectively control the patterns of amino acid substitutions. Averaged over time and across proteins, these evolutionary pressures are sufficiently consistent to produce global substitution patterns that can be used to successfully find homologues, infer phylogenies, and reconstruct ancestral sequences. Although the factors which govern the variation of protein substitution rates has received much attention, the influence of thermodynamic stability constraints remains unresolved. Here we develop a simple model to calculate amino acid rate matrices from evolutionary dynamics controlled by a fitness function that reports on the thermodynamic effects of amino acid mutations in protein structures. This hybrid biophysical and evolutionary model accounts for nucleotide transition/transversion rate bias, multi-nucleotide codon changes, the number of codons per amino acid, and thermodynamic protein stability. We find that our theoretical model accurately recapitulates the complex pattern of empirical rates observed in common global amino acid substitution matrices used in phylogenetics. These results suggest that selection for thermodynamically stable proteins, coupled with nucleotide mutation bias filtered by the structure of the genetic code, is the primary global driver behind the amino acid substitution patterns observed in proteins throughout the tree of life.

## Introduction

Protein amino acid sequences change due to spontaneous mutations at the DNA level. Amino acid exchange rates depend not only on the background mutation rate, but also on how the mutation at the protein level impacts overall organismal fitness. Empirical models that describe amino acid substitution frequencies have wide-ranging applications in bioinformatics and evolutionary science, including homology search, phylogenetics, ancestral sequence reconstruction, and the prediction of functional residues. While such empirical models have tremendous practical utility, the importance of various fitness pressures in shaping the patterns of amino acid substitutions is still unknown. To answer this question, we need biophysical models that predict amino acid substitution rates from first principles. Competing evolutionary hypotheses could then be evaluated and tested by comparison with empirically measured substitution rates. Amino acid mutations can impact fitness by modifying protein properties such as catalysis, binding, expression, aggregation, non-specific interactions, and protein stability, the latter of which is known to be a major selection pressure at most sites for all proteins (Bloom and Arnold 2009; Wylie and Shakhnovich 2011; Araya et al. 2012; Jacquier et al. 2013). Here we develop a model from first principles that predicts amino acid substitution rates by assuming that the fitness effects of amino acid mutations arise from changes in protein stability alone. By comparison against empirical rate data, we evaluate the ability of thermodynamic stability to explain global amino acid substitution patterns.

Phylogenetic models of protein evolution explicitly describe amino acid substitution processes using instantaneous rate matrices whose parameters are inferred from observed sequences using statistical methods. These models describe the substitution pattern as an aggregated Markov process in which sites evolve independently and with a substitution rate that depends on the identity of the current amino acid (e.g., the WAG replacement matrix (Whelan and Goldman 2001)). More recent models also incorporate rate variation across sites (e.g., the LG replacement matrix (Le and Gascuel 2008)). To understand the origin of the variation in amino acid substitution rates, several studies have established correlations between empirical rate variation and biophysical amino acid descriptors (such as hydrophobicity, secondary structure propensity, charge, codon count, and codon table structure) (Tomii and Kanehisa 1996; Venkatarajan and Braun 2001; Atchley et al. 2005; Creixell et al. 2012). While these phenomenological correlations help identify factors that influence amino acid substitution rates, biophysical models are required for a full mechanistic understanding of the underlying fitness constraints that cause the empirical substitution patterns.

Over the past two decades, mechanistic models that combine evolutionary dynamics and protein biophysics have led to significant advances in our understanding of the causes of evolutionary rate variation in proteins (Liberles et al. 2012; Sikosek and Chan 2014; Echave et al. 2016; Echave and Wilke 2017). Biophysical models of rate variation typically treat molecular fitness as dominated by fold stability or the stability of an active structure. Evolutionary trajectories simulated with stability fitness models have shown how fluctuations in the structural environment within proteins influence substitution rates (Pollock et al. 2012) and produce epistatic effects between sites (Shah et al. 2015). However, such thermodynamic evolutionary models have yet to be applied to understand the origin of amino acid substitution patterns.

Here we develop a biophysical model of amino acid substitution rate variation to investigate the evolutionary and thermodynamic basis for substitution patterns in proteins. We first calculate the thermodynamic effects of amino acid mutations in native protein structures and combine a fitness function solely based on protein stability (Paul D Williams et al. 2006) with a position-specific codon-level model of sequence evolution (Halpern and Bruno 1998) to predict global amino acid substitution rates. Using this model, we find that our calculated amino acid substitution rates are strongly correlated with experimental values described by global empirical rate matrices such as the widely used LG matrix. Most of the empirical amino acid substitution pattern can be explained wholly by mutation combined with selection pressure to maintain thermodynamically stable protein structures.

## Results

### A parametric all-atom thermodynamic model of protein evolution

We selected a non-redundant, curated set of 52 high-quality protein structures for the basis of our analysis (Kumar 2006; Kellogg et al. 2011). These 52 protein structures were subjected to a computational analog of saturated mutagenesis. For each site in every protein, we calculated the thermodynamic effect (ΔΔG) of all possible amino acid mutations with the Rosetta macromolecular modeling suite (Leaver-fay et al. 2011). We assume that fitness is proportional to the fraction of folded protein (Paul D Williams et al. 2006), which is determined by the *ΔΔG* value, and then calculate the probability of fixation in a finite population using the Kimura equation (Kimura 1957; Kimura 1962) (Fig. 1). A global amino acid substitution matrix can then be constructed from these amino acid fixation probabilities and a codon-level model of nucleotide mutation. There are only four free parameters in this mechanistic evolutionary model: (1) the free energy of the native protein, *ΔG*_*nat*_, (2) the effective population size, *N*_*e*_, (3) the nucleotide transition rate, *κ*, and (4) a whole-codon rate parameter, *ρ*. Values of these four parameters are required to calculate a substitution matrix; because we do not know the values of these parameters *a priori*, we optimize them with the method of maximum likelihood. The end-result of this method is an amino-acid replacement matrix (189 free exchangeability rates) and an amino acid equilibrium frequency vector (19 free probabilities) that has been fit using only four parameters, each with a clear physical and evolutionary interpretation.

**Fig. 1.**
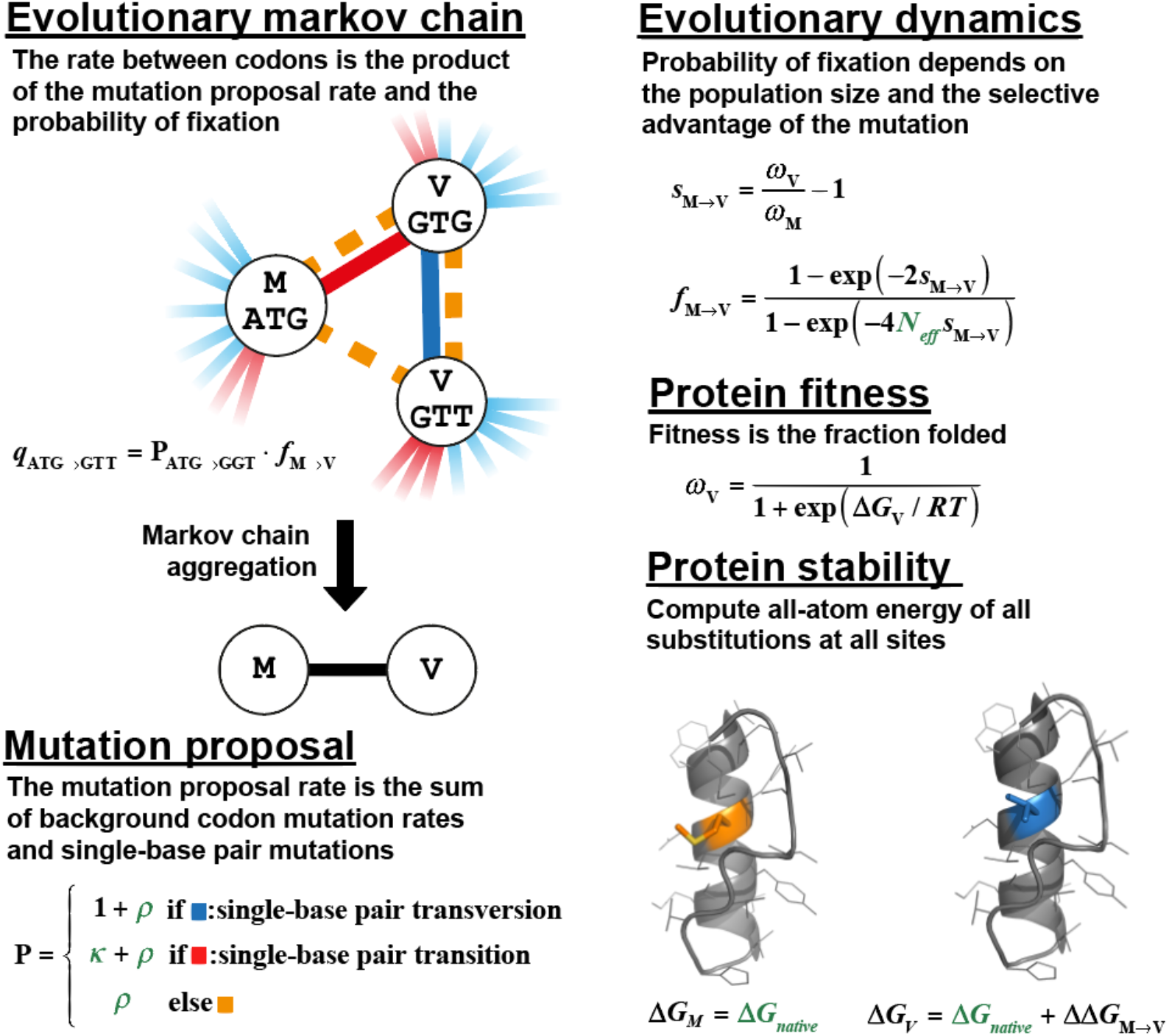
Overview of method illustrated for a M to V mutation. The rate of substitution between any two amino acids in a protein is evaluated based on an underlying codon mutation Markov process. Mutations arise according to the mutation proposal model and fix depending on the fitness difference according to an evolutionary dynamics model. Fitness is proportional to the fraction folded (i.e., the probability of the native state) based on an all-atom energy function. The model contains four free parameters (*ΔG*_*nat*_, *N*_*e*_, *κ*, *ρ*) that are simultaneously optimized.

### Thermodynamic effects of amino acid mutations

A full atomistic model of protein structure and energetics is necessary to realistically model site-specific behavior. However, computing the folding stability of proteins at this level of detail is currently intractable for simulations of protein evolution, as it requires extensive sampling of alternative conformations for every sequence evaluated. Instead, we treat the folding stability of native sequences (*ΔG*_*nat*_) as a global free parameter in the model. In contrast to folding stability, the free energy change upon amino acid mutation (*ΔΔG*) is both reasonably fast to compute and fairly accurate (r^2^=0.56 between prediction and experiment (Park et al. 2016)). We therefore calculate the folding stability of a sequence variant at site *L* as:

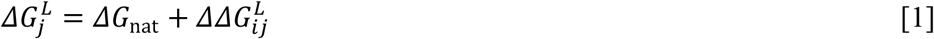

where *i* indicates the index of the native amino acid at site *L* and *j* indicates the index of the proposed amino acid mutation at that site. The equilibrium properties of each position are conditioned only on the fitness pressure exerted by the native structural environment in the selected proteins. In contrast to using a mutation-after-mutation simulation method, this eliminates the need for sequence pre-equilibration, avoids compounding errors of model and energy function, and allows computation of a site-specific Q-matrix from just 20 ΔΔG calculations (Halpern and Bruno 1998).

### Protein fitness as a function solely of thermodynamic stability

To understand to what degree thermodynamic stability explains the process of protein evolution, we base our fitness function on folding stability alone. Following previous work (Paul D. Williams et al. 2006), we assume that a protein’s contribution to fitness is proportional to the fraction of the protein folded into its functional conformation. The fraction of protein folded ω for a two-state folding model with stability 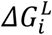 is given by

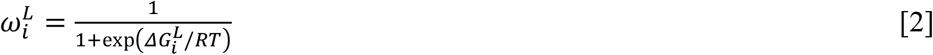

### Fixation probability in a finite population

Given enough time, a proposed mutation will either spread throughout the population and fix or be purged due to negative selection or genetic drift. We assume throughout a monomorphic population evolving under weak-mutation (Golding and Felsenstein 1990). The probability of fixation when amino acid *i* mutates to amino acid *j* at site *L* depends on the relative change in fitness due to that mutation (the selection coefficient 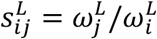) and the effective population size *N*_*e*_ (Kimura 1962):

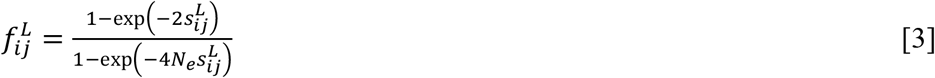

The effective population size *N*_*e*_ is another global free parameter in the model. The fixation probability we use above is for diploid populations with non-overlapping generations in which a new mutation arises as a single copy. Similar equations, with exponential terms differing by a factor of two, apply to haploids and overlapping generations (Ewens 1979), but the effects on our analysis are negligible for large population sizes (e.g., *N*_*e*_ > 100).

### Nucleotide and codon mutation proposals

Most spontaneous mutations involve changes to single-base pairs. Due to the intrinsic chemical properties of nucleotides (M. D. Topal and Fresco 1976; M.D. Topal and Fresco 1976), point mutations occur with greater rates for transitions (pyrimidine to pyrimidine or purine to purine) than transversions. To capture this bias in our model, the transversion rate is fixed at 1.0 and the transition rate *κ* is treated as a free parameter. A smaller subset of mutations involve changes of multi-nucleotides or whole codons, for instance as may result from insertions, deletions, or tandem mutations due to error-prone replication (Harris and Nielsen 2014) or UV damage (Reid et al. 1993). These types of mutations have not been modeled previously in structure-based simulations of evolution, but they play an important role in the evolution of natural sequences (Kosiol et al. 2007). We model the rate of whole-codon mutations with a single parameter *ρ*. The mutation proposal probability *P* at a given site is:

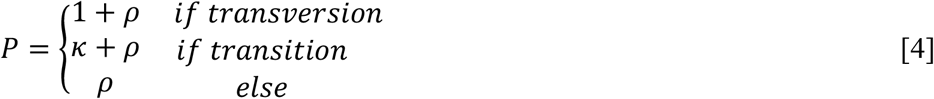

### Construction of amino acid rate matrices from codon mutations and protein fixation

We model the site-specific evolutionary process as a continuous-time Markov process, where the unit of change is the codon and the relative instantaneous substitution rate 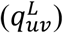 from codon *u* with amino acid *i* to codon *v* with amino acid *j* is the product of the mutation proposal rate, *P*_*uv*_, and the fixation probability, 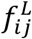 (Halpern and Bruno 1998):

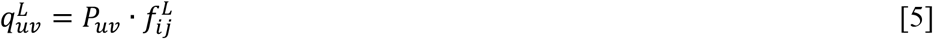

Codons representing the same amino acids are then aggregated, allowing determination of site-specific amino acid flux matrices. The flux matrices are averaged across sites and proteins to construct a global 20×20 amino acid instantaneous rate matrix *Q*. For comparison with empirical global substitution matrices such as JTT, WAG, and LG, and for use in phylogenetic analyses, the *Q*-matrix was decomposed into a diagonal amino acid equilibrium probability matrix, *π*, and an independent symmetric exchangeability matrix *R* (Le and Gascuel 2008):

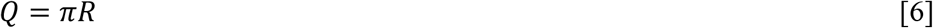

The equilibrium probabilities describe the expected frequency distribution of amino acids when the substitution process has progressed long enough that it reaches equilibrium.

### Optimized model parameters provide typical values

Using the computed mutation energies (ΔΔG), an optimal exchangeability matrix *R* was found by optimizing values of the four parameters of the model *θ* = {*ΔG*_*nat*_, *N*_*e*_, *κ*, *ρ*} by maximizing the phylogenetic likelihood for the 52 non-redundant protein families. With grid-search optimization, we found an optimal parameter set of *ΔG*_*nat*_=−6.0±0.1 kcal/mol, log(*N*_*e*_)=3.8±0.1, *κ*=2.1±0.1, and *ρ*=0.11±0.01 (standard error estimated by bootstrap (Efron 1979), see supporting methods). We refer to this optimized matrix as the Thermodynamic Mutation-Selection (TMS) matrix. Because the TMS model parameters are based on fundamental quantities in population genetics (*N*_*e*_), spontaneous mutation processes (*κ*, *ρ*) and protein thermodynamics (*ΔG*_*nat*_), the optimal parameter values can be compared with independent empirical measurements from protein biochemistry and population genetics. As discussed further below, these optimal values from our thermodynamic, evolutionary model correspond surprisingly well to representative physical and biological values.

The stability of natural proteins is typically between −5 to −10 kcal/mol (Ghosh and Dill 2010), while effective population sizes vary over many orders of magnitude. For instance, the effective population size for humans is on the order of 10^4^ (Yu et al. 2004), while *E. coli* have an population size of approximately 10^7^ (Charlesworth and Eyre-Walker 2006). The optimal *ΔG*_*nat*_ and *N*_*e*_ TMS values compare well with these ranges. By inspecting the likelihood surface (see Fig. S1), the stability *ΔG*_*nat*_ and population *N*_*e*_ parameters are seen to be highly correlated, with optimal parameters found along a line given by

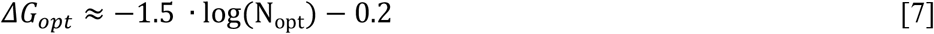

A similar dependence of protein stability *ΔG* on *N*_*e*_ has been noted previously (Wylie and Shakhnovich 2011). The strong correlation between *ΔG* and *N*_*e*_ suggests that they represent a single latent parameter, reducing the effective parameter space of our model from four to three.

In contrast to stability and population size, the parameters describing the mutation process (*κ* and *ρ*) are largely independent (Fig. S1). The optimal transition/transversion bias *κ*=2.1 is consistent with experimental data for spontaneous mutation rates from *E. coli* (Lee et al. 2012), where *κ*≃2.6 (assuming a K80-type Markov model (Kimura 1980), see supplementary). For the multi-nucleotide mutation parameter *ρ*, the MLE is 0.1. Obtaining direct empirical estimates of *ρ* is difficult, but for indels of less than four nucleotides in *E. coli*, the spontaneous mutation rate is approximately 10% of the single-nucleotide mutation rate (Lee et al. 2012). In eukaryotes, multi-nucleotide mutations comprise approximately 3% of all mutations (Schrider et al. 2011) and have been shown to be important in phylogenetic tests (Venkat et al. 2018). Given that processes other than indels can result in multi-nucleotide mutations, and that indels typically result in multiple codon mutations, our *ρ*_*opt*_ value appears reasonable.

### The thermodynamic TMS model reproduces empirical substitution patterns

The majority of the variation in empirical amino acid substitution matrices can be explained by our thermodynamic evolutionary model, as judged by the logarithmic correlation coefficient with the ML TMS matrix (Fig 2). The TMS matrix appears to be a rather typical exchangeability matrix, as the average correlation of TMS with many widely used empirical matrices is r^2^=0.54, whereas the correlation of those empirical matrices with each other (excluding TMS) is r^2^=0.52 (Fig 2). The average correlation with globular matrices is higher (r^2^=0.59), whereas the correlation with mitochondrial matrices is substantially lower (r^2^=0.42). For example, the TMS exchangeability matrix has an r^2^ of 0.67 and 0.64 with the widely used WAG (Fig. S2) and LG substitution matrices (Fig. 3), respectively. The lower correlation with mitochondrial matrices is likely due to the preponderance of transmembrane proteins in the mitochondrial datasets (Jimenez-Morales and Liang 2011; Le et al. 2017), as the Rosetta energy function used in our thermodynamic free energy calculations is intended for soluble proteins. Breaking exchangeabilities down by individual amino acids, we see the lowest correlation for cysteine and proline (r^2^=0.50), likely reflecting shortcomings of the energy function and limited modeling of backbone flexibility. The other amino acids have higher correlations, up to r^2^=0.85 (Fig. 4).

**Fig. 2.**
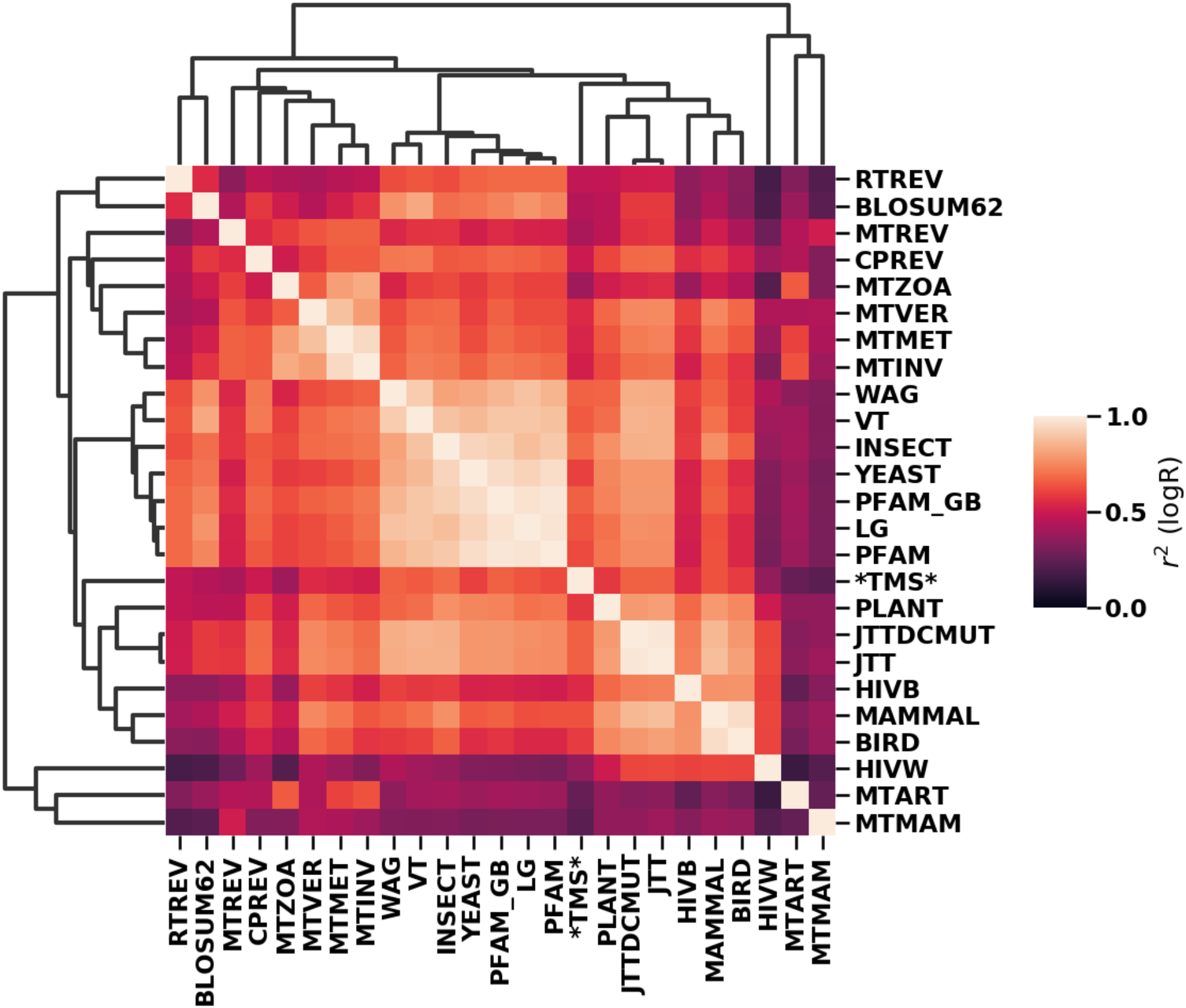
Heatmap of correlations between popular exchangeability matrices. Squared Pearson correlations (r^2^) were computed in log-space over the 190 exchangeability parameters. Matrices were clustered based on r^2^ using hierarchically clustering (Virtanen et al. 2020).

**Fig. 3.**
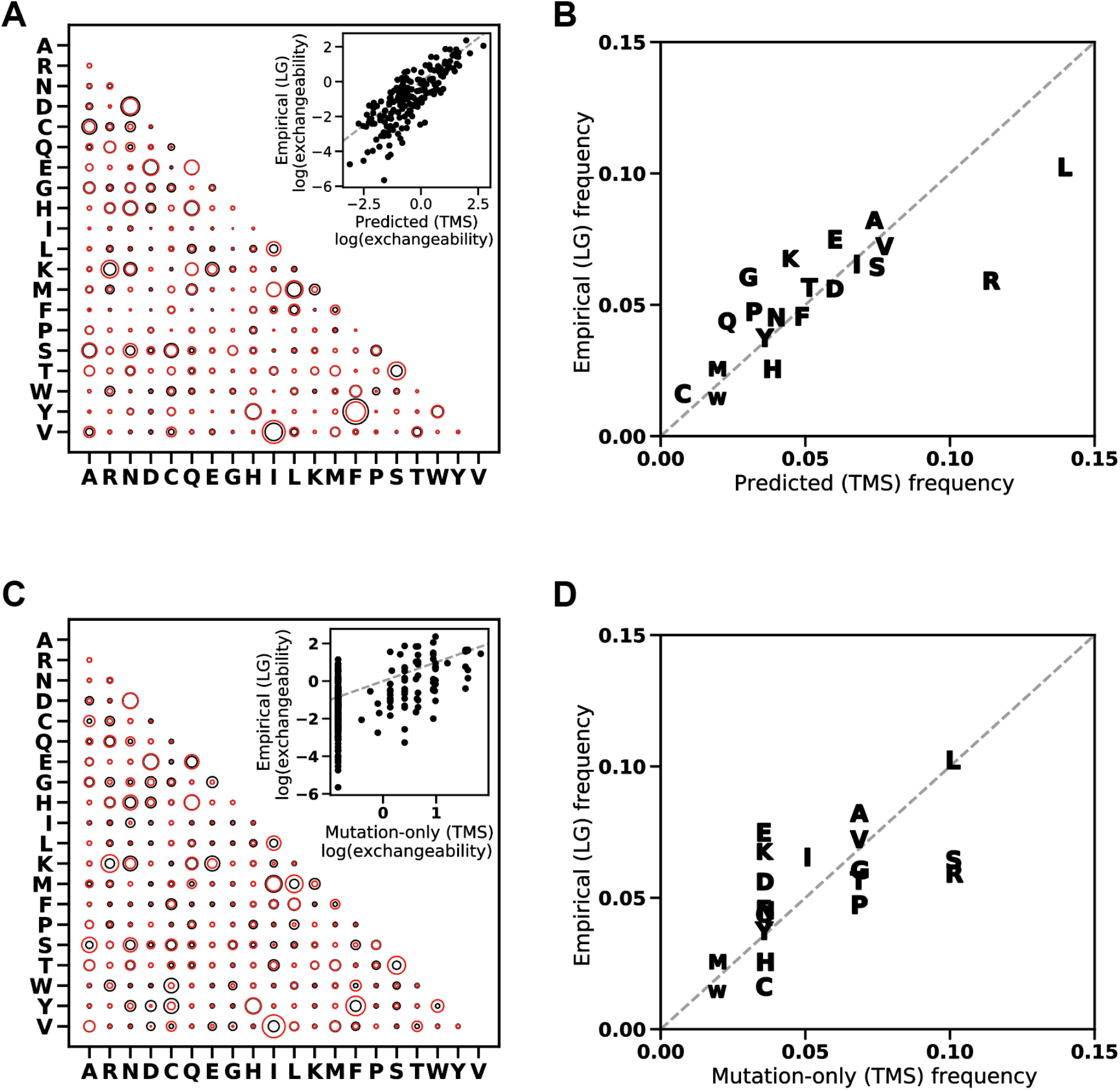
The TMS model recapitulates the mean substitution behavior between amino acids. A) Comparison of the amino acid exchangeability matrix predicted by TMS (black circles) and a phylogenetic empirical exchangeability matrix (LG, red circles). Inset, correlation between exchangeabilities from TMS (*x*-axis) vs LG (*y*-axis). B) Correlation between amino acid equilibrium probabilities predicted by TMS (*x*-axis) and values from LG (*y*-axis). Identity shown as dashed line. C) The mutation-only exchangeability matrix, without selection for stability, vs LG. D) Correlation between mutation-only equilibrium probabilities and those from LG. Identity shown as dashed line.

**Fig. 4.**
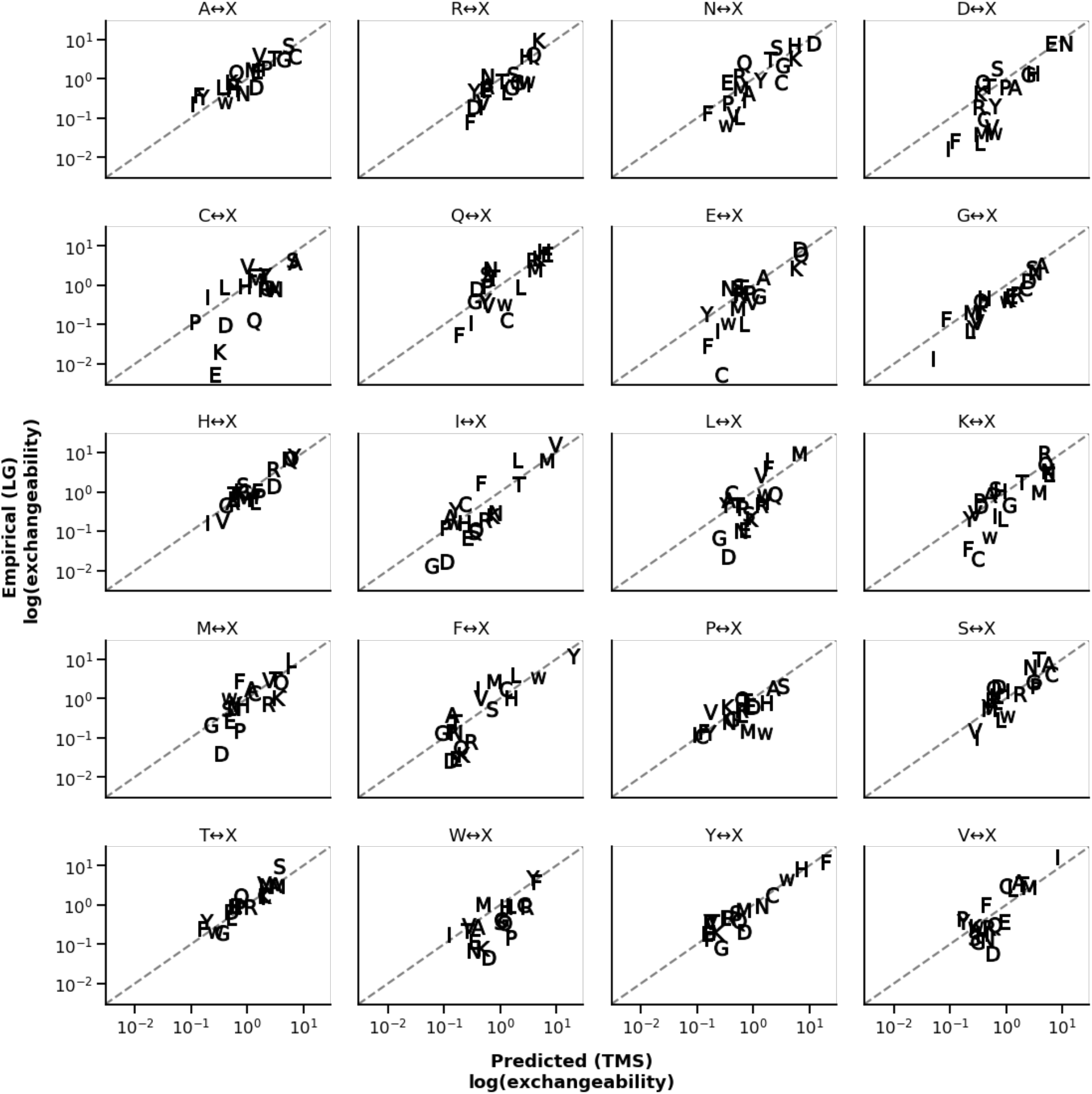
The TMS model recapitulates amino acid specific exchangeabilities. Identity is shown as a dashed line.

The high correlation between TMS and other experimental matrices like LG is a result of both the thermodynamic model, in which more chemically similar amino acids have higher fixation probabilities, and the mutation model, which biases amino acid replacements in the genetic code due to preferred nucleotide mutations, codon number, and codon connectivity. To understand the impact of the genetic component on the correlations, we computed a “mutation-only” *R*-matrix, assuming only the connectivity of the codon table and the transition-transversion bias by setting all fixation probabilities in eq. 5 to 1 and setting *ρ*=0.10 and κ=2.1 to their optimal values. This mutation-only *R*-matrix accounts for 32% of the variation in the LG substitution pattern (Fig. 3C), suggesting that genetic and thermodynamic factors contribute roughly equally to the patterns of empirical amino acid replacement rates.

Correct recapitulation of amino acid exchangeabilities is strongly dependent on the thermodynamic component of the model. For example, using LG as the standard for comparison, the empirical exchangeability for tryptophan to tyrosine is underestimated by 7.4-fold by the mutation-only *R*-matrix but is underestimated by 1.1-fold by TMS. Similarly, the mutation-only model overestimates the empirical exchangeability of asparagine to phenylalanine by 4.8-fold, while TMS overestimates it by 1.3-fold.

Our ML TMS matrix was optimized over the exchangeabilities and not the amino acid equilibrium frequencies. Nevertheless, from the decomposition of the Q-matrix (given in eq. 6), we can provide the implied TMS equilibrium frequencies and compare them with those provided with common exchangeability matrices (Fig. S3). For example, the calculated TMS equilibrium frequencies correlate well with those from LG (linear r^2^=0.65), but suggest unmodelled fitness effects. For instance, we do not model disulfide formation, and unsurprisingly the stationary frequency of cysteine calculated from TMS is 2.8-fold too low. Likewise, we ignore codon usage biases, which could explain the overestimation of the background frequency of arginine and leucine, both of which have exceptionally skewed codon usage (Plotkin and Kudla 2011).

Although our model correlates strongly with empirical exchangeability matrices like LG, some of the variation in the rates remains unexplained. One possible contribution to this discrepancy stems from the fact that we selected a set of proteins that are different from those used to infer the LG-matrix. Our set of proteins will likely have a slightly different substitution process than the LG set. To investigate this possibility, we used a maximum likelihood phylogenetic inference method (Dang et al. 2011), similar to that used for the construction of the LG matrix, to infer exchangeability rates from the sequence alignments for our set of benchmark proteins (Fig. S4). The inferred phylogenetic exchangeability matrix for our proteins is highly correlated with LG (r^2^=0.97), indicating that our benchmark proteins are similar to those that LG was derived from, and that stochastic errors originating from the rate-inference itself are minimal.

A second source of the unexplained variation could be errors in the energy function used for ΔΔG prediction, as Rosetta energies have an imperfect correlation with empirical values (r^2^=0.56). We explored this possibility by simulating noisy data using empirically determined ΔΔG Rosetta prediction errors. This analysis suggested at least 21% of the unexplained variance could be caused by energy function errors (see supplementary). Finally, our mutation-selection model operates at the codon level, producing rates that are subsequently aggregated to the amino acid level – a reduction in dimensionality that can affect the assumed Markov property of codon and amino acid evolution (Spielman and Shapiro 2020). It is possible that our codon-level model could explain codon exchangeability matrices better than amino acid matrices. Additionally, when determining an amino acid exchangeability matrix like LG by phylogenetic methods, all 189 exchangeability rates are free parameters that are estimated independently from the sequence data. In contrast, our codon model is highly constrained by only two parameters (*κ* and *ρ*) and the K80 assumption of equal nucleotide frequencies. More complex mutational models could provide higher correlations at the expense of model simplicity and interpretability.

### The thermodynamic model explains some trees better than empirical substitution matrices

The purpose of our study is to see how far a simple thermodynamic and mutation-selection model could go in explaining empirical amino acid substitution rates; it was not intended to provide a better substitution matrix for practical use. Nevertheless, it is interesting to see how well our TMS matrix fares in phylogenetic analyses. For each of the 52 individual phylogenetic trees in our benchmark set, we compared the TMS maximum likelihood values to the LG maximum likelihood values for the same alignments. The mean TMS likelihood was −13664 while the mean LG likelihood was −13492, a difference of 172 on average in favor of LG. For most trees the LG matrix is better, but for 3 of the 52 trees TMS has a higher likelihood than LG. Using a larger set of 500 Pfam alignments (Yang 1994; Le and Gascuel 2008), independent of both our model parameters and LG, we similarly found that in 8 alignments TMS had higher likelihood than LG (see supplementary table 1). Thus, for a certain subset of protein families, TMS does appear to provide a better substitution model than LG.

## Conclusions

We present an approach to predict relative rates of amino acid substitutions in evolution from first principles by combining population genetics and protein biophysics. Our model is based on an extremely simple assumption: the fitness of a gene variant is controlled only by constraints on protein stability. Nevertheless, this naive model can remarkably recapitulate the complex amino acid substitution patterns seen in empirically derived rate matrices. Our model incorporates transition/transversion mutation biases, multi-nucleotide codon changes, variation in codon counts, and detailed atomic interactions in proteins. By calculating the fitness effects of mutations in the structural environment of native sequences, this method can identify biologically and physically meaningful model parameters that are otherwise difficult to estimate directly from protein sequence datasets. We find that most of the substitution behavior of amino acids in proteins can be explained by a simple evolutionary fitness model that captures (1) the thermodynamic effects of mutations, (2) the biases in spontaneous mutations, and (3) the structure of the genetic code.

## Methods

### Protein dataset

We selected a non-redundant subset of 52 proteins structures from a curated set of high-quality protein structures (Kumar 2006; Kellogg et al. 2011). All structural modeling was done with the Rosetta macromolecular modeling suite (Leaver-fay et al. 2011). To ensure structural diversity, sequence redundancy was decreased so that no sequence shared more than 60% identical positions with any other sequence. Before analysis, each structure was adapted to the energy function using the *FastRelax* protocols as described by Nivon et al. (Nivón et al. 2011) allowing cartesian space minimization.

### Prediction of the energetic effects of amino acid mutations

The ΔΔG prediction method is based on a modified version of the method presented by Park et al. (Park et al. 2016), but with a cutoff in the Lennard-Jones potential set to 6.0 Å. This ΔΔG method samples backbone degrees of freedom for the mutated and neighboring residues in the sequence and allows repacking of all-side chains in energetic contact (>0.1 kcal/mol) with the mutated residue.

### Markov chain aggregation and averaging

To calculate the amino acid substitution rate matrix 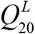 for site *L* from the codon substitution rate matrix 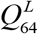, we first determine the frequency of each codon (Iwasa 1988; Sella and Hirsh 2005):

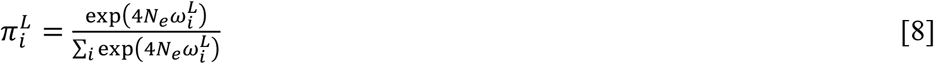

Next, we determine the rate between two amino acids (*i*, *j*) with codons *u* ∈*i* and *v* ∈ *j* using the aggregation approach presented by Yang et al. (Yang et al. 1998):

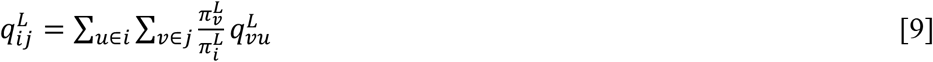

where 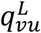 is the rate between codon *v* to *u* at site *L*. The flux between a pair of amino acids at a site *L* is:

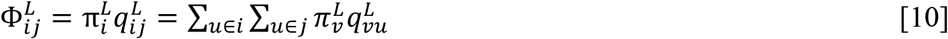

The site-specific rate *μ*^*L*^ (the total flux) is:

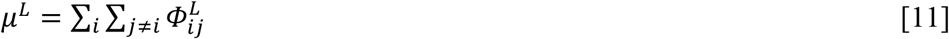

We normalize the site-specific flux matrix, 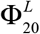, so that a unit of time corresponds to one expected amino acid substitution per site.

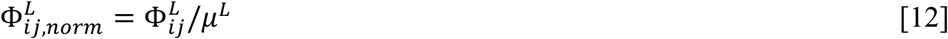

To calculate the mean instantaneous rate for amino acid *i* to *j* we average over sites by normalizing by the equilibrium frequency of an amino acid *i* at that site:

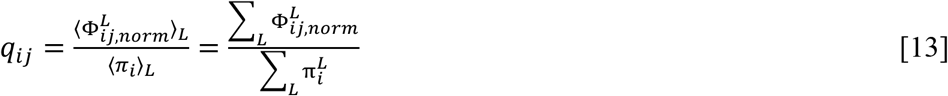

Sites with rates with μ<10^−10^ were excluded from the analysis to avoid numerical errors. Next, we determine exchangeability matrix *R*_20_ = (*r*_*ij*_) as

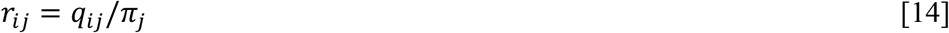

### Model parameterization based on phylogenetic trees

Optimal values for the four free parameters of the model 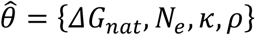 were determined by maximizing the sum log-likelihood of phylogenetic trees for the 52 non-redundant protein alignments with a total of 8907 sites:

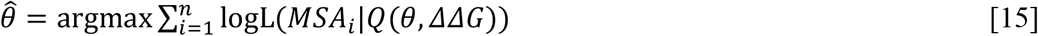

The maximum likelihood trees (optimizing branch lengths, topology, equilibrium frequencies, and site rate variation parameter α) were determined with IQ-TREE (model TMS+FO+G4, where TMS is an exchangeability matrix calculated from specific values of *ΔG*_*nat*_ = −5.91, *N*_*e*_ = 10^3.8^, *κ* = 2.1, *ρ* = 0.1). (Minh et al. 2020). Note that because we use the IQ-TREE “FO” model option, rather than specifying the equilibrium frequencies predicted by our TMS model, this method optimizes over the TMS exchangeabilities alone. To speed the calculation the tree search was seeded with a tree inferred using LG. To find the parameterization that maximizes the sum log-likelihood, we performed a grid-search over the parameter space. Grid search optimization was performed over linearly spaced steps in the ranges 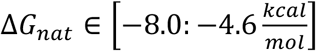, *κ* ∈ [1.4: 2.8] and over log-linearly spaced steps in the ranges *ρ* ∈ [0.05: 0.25], *N* ∈ [10^3^: 10^5^]. The maximum likelihood TMS matrix is available in supplementary materials.

### Correlations between substitution matrices

The agreement between exchangeability matrices was quantified using a Pearson correlation coefficient calculated in log-space for the 190 exchangeability parameters.

### Data availability

Data, substitution matrices, and scripts for analysis are available here: https://github.com/Andre-lab/TMS

## Acknowledgment

CN and IA was supported by a grant from the Swedish Research Council (2015-04203). DLT was supported by NIH grants R01GM096053 and R01GM132499.

## Supplementary information

## Supplementary methods

### Estimating transition/transversion bias in E. coli

Lee et al. (Lee et al. 2012) reported the counting data of spontaneous nucleotide mutations in *E. coli*. The data in their table 3 is the mutation flux. The mutation flux is a function of the instantaneous mutation rates and the equilibrium frequencies: *Φ*_*ij*_ = *q*_*ij*_ ∙ *π*_*ij*_. In accordance K80-type Markov model (Kimura 1980), we assume that the equilibrium frequency of the nucleotides is 0.25 for all and calculate the transition/transversion rate ratio (κ):

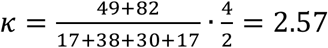

### The protein misfolding-fitness function behaves as the sigmoidal stability-fitness function with a modified offset

Another fitness model was proposed by Drummond and Wilke (Drummond and Wilke 2008). They found that a major selection pressure is selection against cytotoxicity caused by protein misfolding. In their model the fraction of unfolded protein has an exponentially decreasing effect on fitness, which depends on a toxicity parameter c and the protein abundance parameter A.

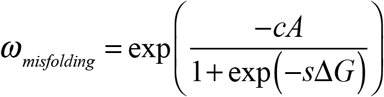

When sΔG is low (sΔG<-3) the misfolding fitness function can be simplified to the fraction folded fitness function although offset by log(cA):

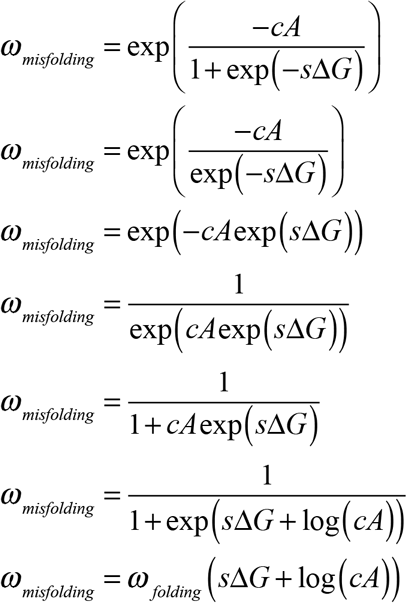

Where ω_folding_ is the fitness function used throughout the paper:

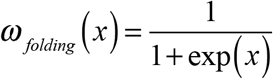

### Estimation of parameter uncertainty by bootstrap analysis

To estimate the uncertainty of the TSM parameters we performed a bootstrap analysis. Specifically, we compute tree log-likelihood (IQ-TREE) for each parameter combination explored in Fig. S1 for each of the 52 sequence alignments individually. Next, using random sampling with replacement (sample size = 52) over the alignment dataset, we took 10.000 bootstrap samples, for each finding the argmax parameter set:

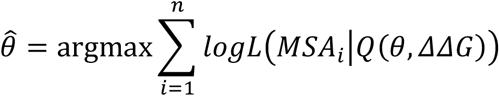

From these samples, we computed the standard deviation over each parameter.

### Estimation of Q-matrix estimation error upon error injection

Deviations between inferred rates and rates predicted with the TSM-model could also originate from ΔΔG prediction. The magnitudes of ΔΔG prediction errors were estimated from the correlation between predicted ΔΔG values from Rosetta and known experimental values for the same proteins and substitutions (σ_err_=1.7 kcal/mol based on 590 measurements). To estimate the impact of such errors, we predicted rates with the TSM model as described in the main text but modified the ΔΔG values by adding errors sampled from the empirical error distribution. The expected correlation between Q-matrix parameters with and without sampled error is r_π+R_^2^=0.79. Thus, ΔΔG prediction error alone likely contributes at least 21% of the variance in the LG matrix that is unexplained by our model. We note that the true error could be greater, as half of the substitutions in the experimental ΔΔG dataset are to alanine, and alanine substitutions are more accurately predicted than other mutations (σ_err,X->A_=1.4 vs σ_err,X->!A_=1.9 kcal/mol).

## Supplementary figures

**Fig. S1.**
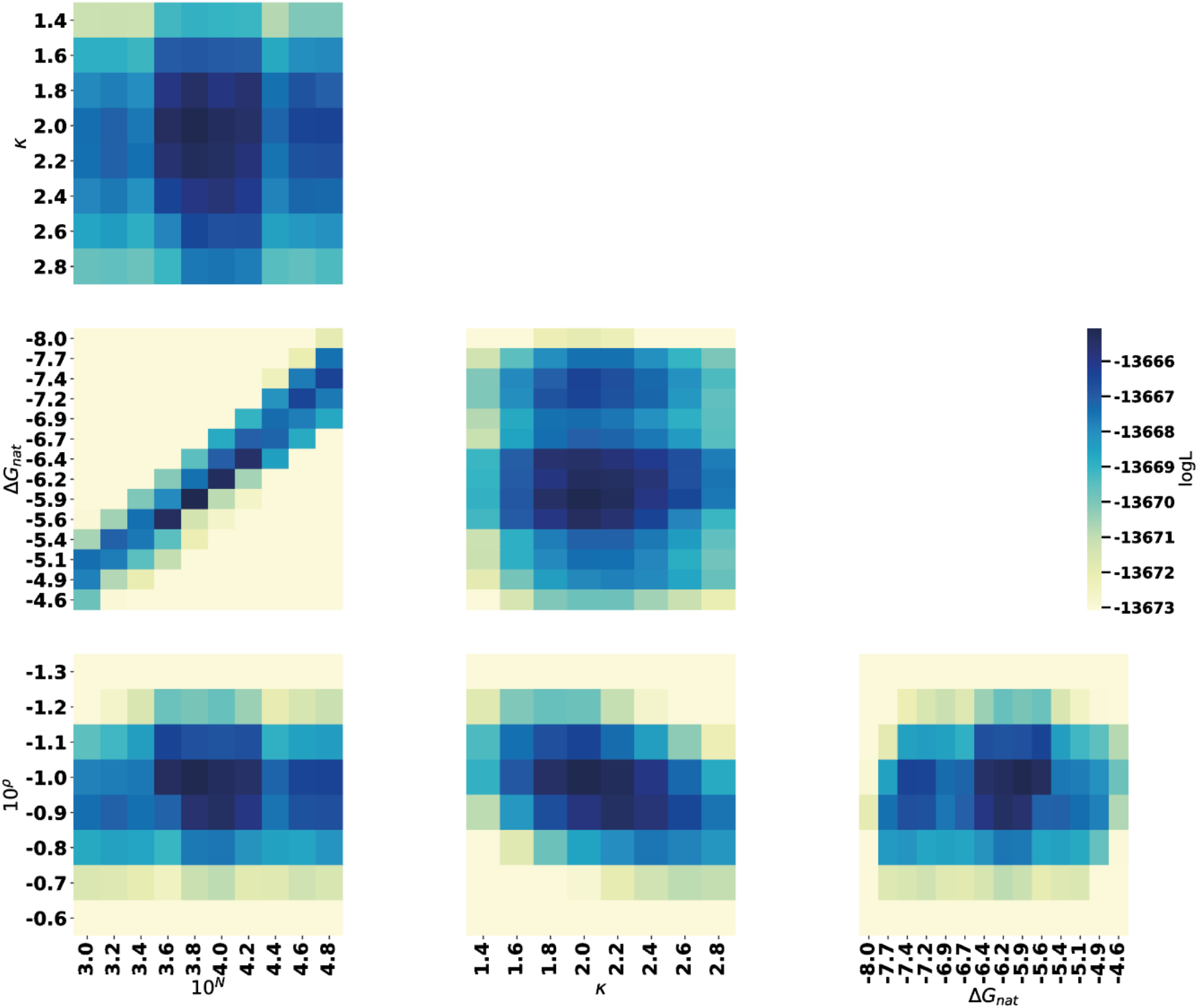
Coupling and sensitivity-analysis of optimal parameter set for the TSM-model. Parameter vs parameter and best MLE in that parameter category.

**Fig. S2.**
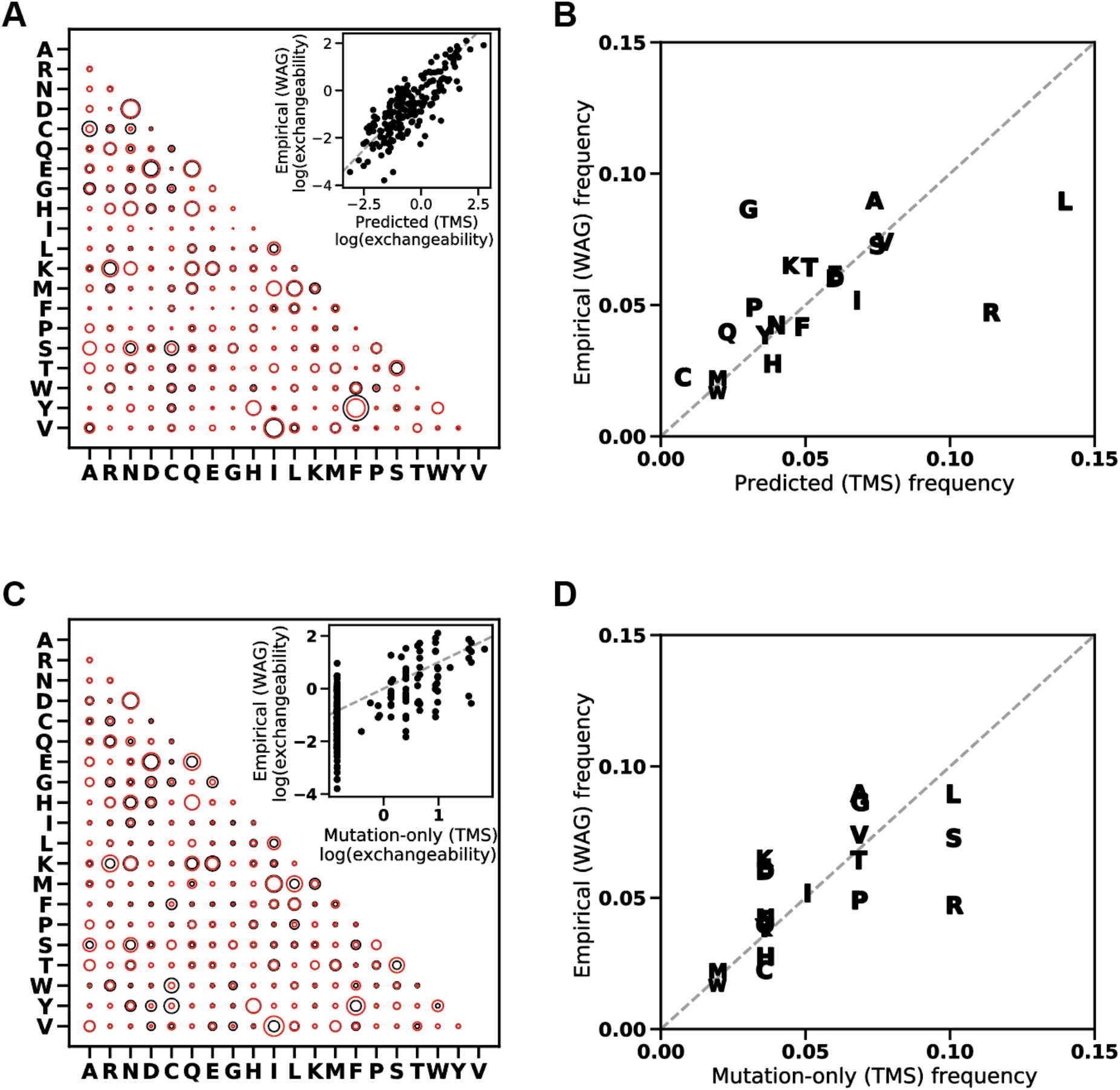
The TMS model recapitulates the mean substitution behavior between amino acids. A) Comparison of the amino acid exchangeability matrix predicted by TSM (black circles) and an empirical sequence-based exchangeability matrix (WAG, red circles). Inset, correlation between exchangeabilities from TSM (*x*-axis) vs WAG (*y*-axis). B) Correlation between amino acid equilibrium probabilities predicted by TSM (x-axis) and values from WAG (y-axis). C) The same as A, but without selection. D) the same as B but without selection.

**Fig. S3.**
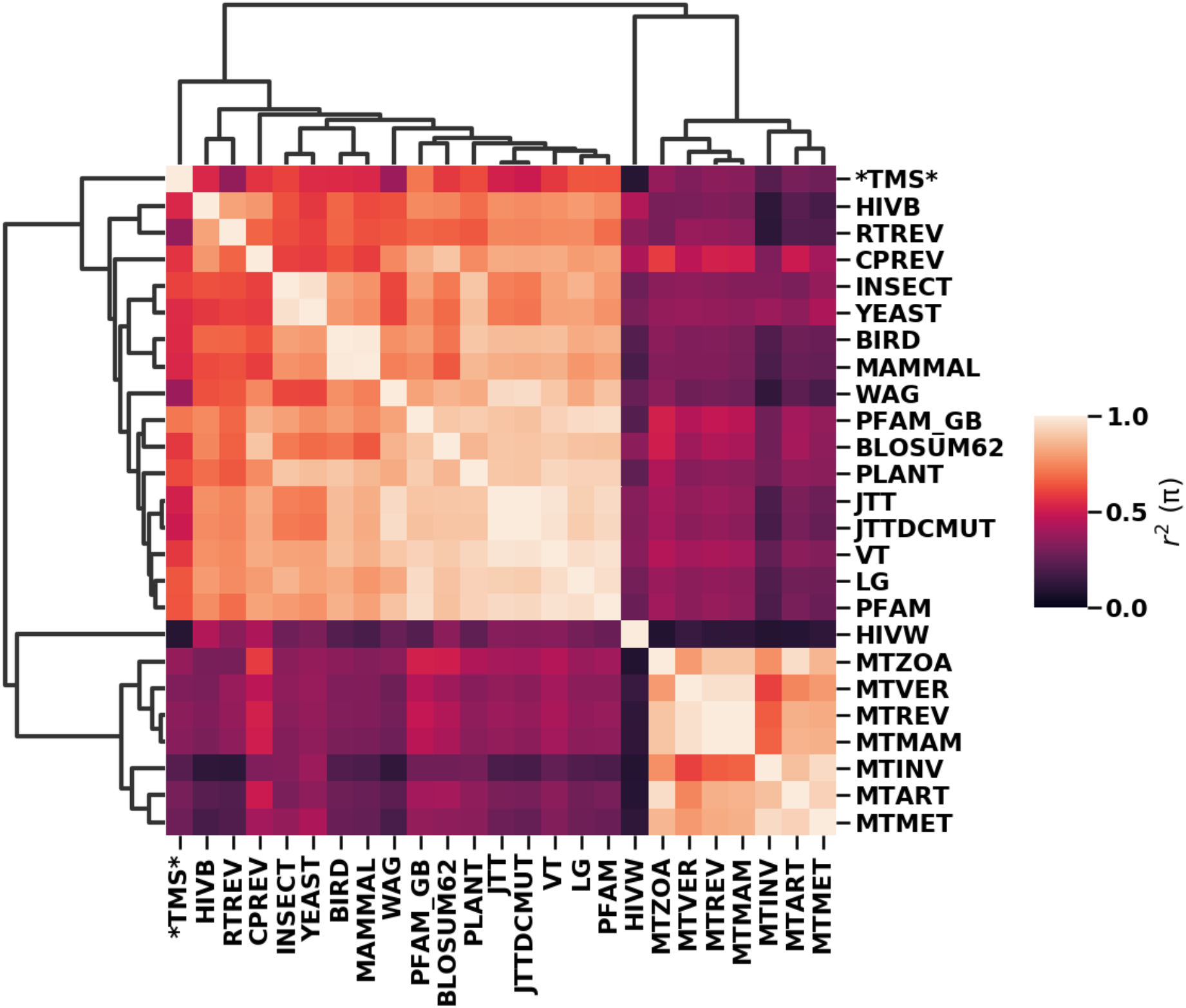
Heatmap of correlations between stationary amino acid frequencies for popular substitution matrices. Squared Pearson correlations (r^2^) were computed in log-space over the 190 exchangeability parameters. Matrices were clustered based on r^2^ using hierarchically clustering (Virtanen et al. 2020).

**Fig. S4.**
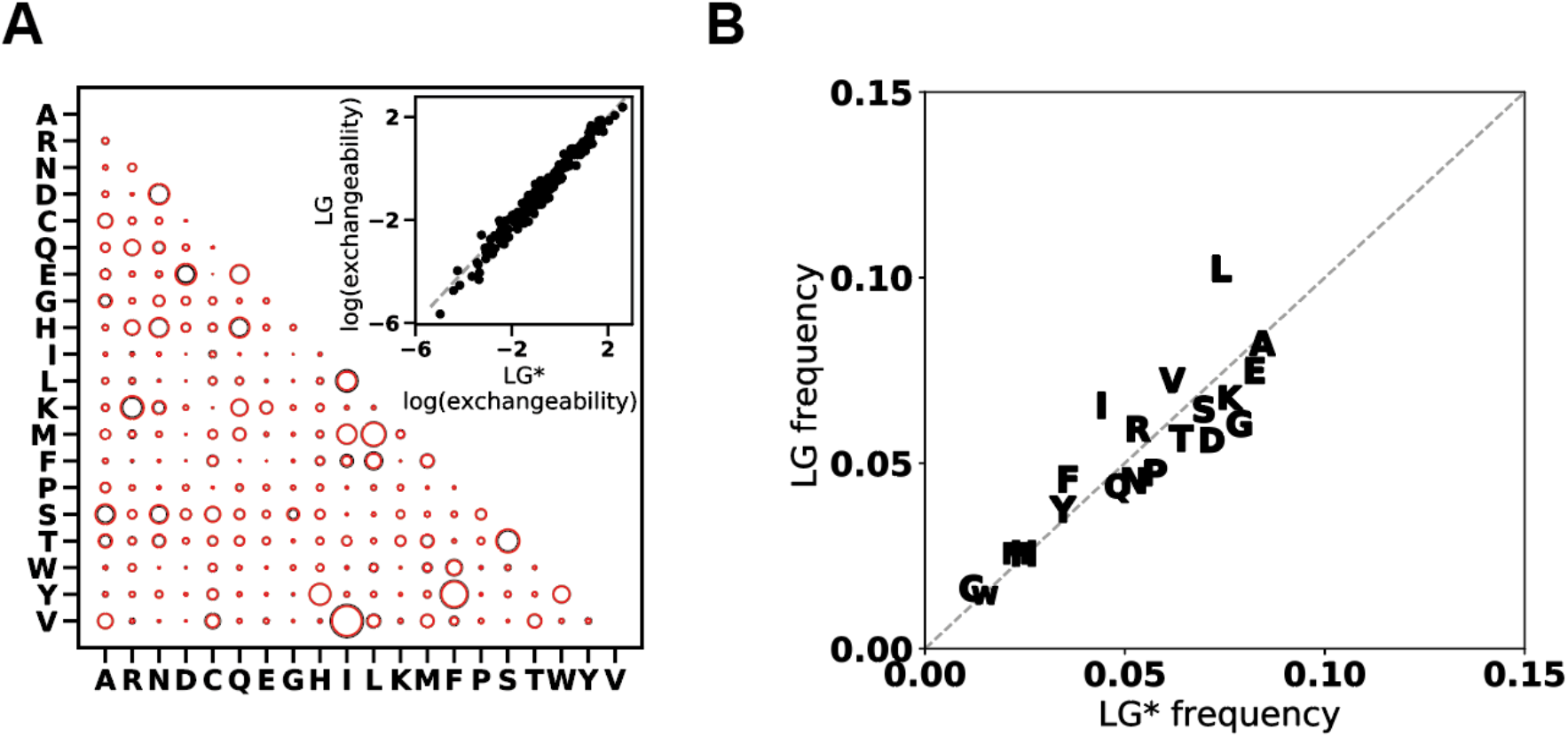
Proteins used for calculating substitution energetics are representative of the alignments used to infer LG. A) Comparison of an amino acid exchangeability matrix inferred from 52 alignments (LG*, red circles) and a state-of-the-art empirical substitution matrix (LG, black circles). Inset, correlation between exchangeabilities. B) Correlation between amino acid equilibrium probabilities inferred from LG* and values from LG. The 52 alignments were pfam seed alignments obtained for proteins for which we calculated substitution energetics.

**Table S1:** Log-likelihoods of trees computed for MSAs in the LG pfam test dataset. The table shows likelihoods for the MSAs with a delta log likehood of less than 5. Out of 500 MSAs, our Q-matrix achieved a higher likelihood than LG for 8.

| **aln** | **TSM logL** | **LG logL** | **ΔlogL** |
| --- | --- | --- | --- |
| Aln2393.txt-gb_phyml | -577.403 | -590.073 | -12.670 |
| Aln6612.txt-gb_phyml | -2493.606 | -2500.485 | -6.879 |
| Aln3459.txt-gb_phyml | -3494.576 | -3499.601 | -5.025 |
| Aln3653.txt-gb_phyml | -1272.407 | -1276.997 | -4.590 |
| Aln5878.txt-gb_phyml | -950.257 | -954.827 | -4.570 |
| Aln1284.txt-gb_phyml | -634.963 | -638.874 | -3.911 |
| Aln3227.txt-gb_phyml | -756.610 | -758.864 | -2.254 |
| Aln3035.txt-gb_phyml | -978.598 | -980.501 | -1.903 |
| Aln4281.txt-gb_phyml | -754.795 | -753.672 | 1.123 |
| Aln6528.txt-gb_phyml | -490.274 | -489.116 | 1.158 |
| Aln5555.txt-gb_phyml | -392.622 | -390.080 | 2.542 |
| Aln2167.txt-gb_phyml | -798.087 | -795.381 | 2.706 |
| Aln1980.txt-gb_phyml | -1019.706 | -1016.945 | 2.761 |
| Aln2041.txt-gb_phyml | -1754.531 | -1751.638 | 2.893 |
| Aln4313.txt-gb_phyml | -1345.046 | -1341.611 | 3.435 |
| Aln4019.txt-gb_phyml | -635.864 | -632.414 | 3.450 |
| Aln1245.txt-gb_phyml | -850.857 | -847.353 | 3.504 |
| Aln0070.txt-gb_phyml | -433.801 | -429.861 | 3.940 |
| Aln4914.txt-gb_phyml | -847.560 | -843.521 | 4.039 |
| Aln2179.txt-gb_phyml | -546.731 | -542.558 | 4.173 |
| Aln4871.txt-gb_phyml | -389.268 | -385.089 | 4.179 |
| Aln2807.txt-gb_phyml | -1758.808 | -1754.351 | 4.457 |
| Aln5867.txt-gb_phyml | -696.083 | -691.502 | 4.581 |
| Aln4013.txt-gb_phyml | -858.934 | -854.133 | 4.801 |

